# Stably Accelerating Stiff Quantitative Systems Pharmacology Models: Continuous-Time Echo State Networks as Implicit Machine Learning

**DOI:** 10.1101/2021.10.10.463808

**Authors:** Ranjan Anantharaman, Anas Abdelrehim, Anand Jain, Avik Pal, Danny Sharp, Utkarsh, Chris Rackauckas

## Abstract

Quantitative systems pharmacology (QsP) may need to change in order to accommodate machine learning (ML), but ML may need to change to work for QsP. Here we investigate the use of neural network surrogates of stiff QsP models. This technique reduces and accelerates QsP models by training ML approximations on simulations. We describe how common neural network methodologies, such as residual neural networks, recurrent neural networks, and physics/biologically-informed neural networks, are fundamentally related to explicit solvers of ordinary differential equations (ODEs). Similar to how explicit ODE solvers are unstable on stiff QsP models, we demonstrate how these ML architectures see similar training instabilities. To address this issue, we showcase methods from scientific machine learning (SciML) which combine techniques from mechanistic modeling with traditional deep learning. We describe the continuous-time echo state network (CTESN) as the implicit analogue of ML architectures and showcase its ability to accurately train and predict on these stiff models where other methods fail. We demonstrate the CTESN’s ability to surrogatize a production QsP model, a >1,000 ODE chemical reaction system from the SBML Biomodels repository, and a reaction-diffusion partial differential equation. We showcase the ability to accelerate QsP simulations by up to 56x against the optimized DifferentialEquations.jl solvers while achieving <5% relative error in all of the examples. This shows how incorporating the numerical properties of QsP methods into ML can improve the intersection, and thus presents a potential method for accelerating repeated calculations such as global sensitivity analysis and virtual populations.

## Introduction

The promises of quantitative systems pharmacology (QsP) are hindered by it’s computational complexity: faster compute speeds will be required for faster drug development [11]. For these large-scale and stiff ordinary differential equation (ODE) models, the compute time required to perform the simulation heavily impacts the speed at which researchers can iterate through model development and perform analyses such as virtual populations and global sensitivity analysis [30]. Given that these aspects have been widely cited as being major time consuming aspects of the preclinical analysis pipeline [5,7,13,14,15,16,17], our goal is to develop a new set of mathematical tools which enable computational scientists to more efficiently and accurately apply model-informed drug development (MIDD) across large complex models.

One promising avenue for generally accelerating modeling and simulation is the incorporating of techniques from machine learning [1,2,19]. Techniques such as surrogates or emulators allow for training machine learning models to capture the behavior of large models in a fast approximation [20]. Such surrogate modeling techniques have demonstrated promise in limited applications in QsP [18]. In this manuscript, we take a deeper look at surrogate modeling and investigate their applicability to the field of QsP models. We will show that while surrogates can be successfully applied to achieve a 5.2x speedup on a large ODE model and a 56x speedup on a large reaction-diffusion partial differential equation (PDE) model, care needs to be taken. We will demonstrate that in many cases it is likely that machine learning techniques will need to be modified in order to be appropriate for this domain.

### Multiscale Models, Stiffness, and the Effect on Machine Learning

One aspect which needs to be addressed with QsP models is their tendency to capture multiple timescale phenomena. An example is how model features like the circadian rhythm or metabolite digestion can span hours while the individual dynamics of important transcription factors can act in microseconds. This timescale separation leads to numerical issues which manifest in highly ill-conditioned Jacobians, a phenomenon which is known as stiffness in the field of ordinary differential equations [21]. Figure 1 is a diagrammatic description of how stiffness causes issues in numerical prediction, where making predictions on the timescale of the slow process cannot be done using local information such as derivatives. This is because such local features change too quickly, giving an inaccurate view of the dynamics of longer term processes. In the space of QsP, this issue normally manifests itself as a problem with ODE solvers: explicit methods like MATLAB’s ode45 are unstable in the presence of these effects, and thus implicit methods like ode15s or CVODE’s BDF methods are required.

**Figure 1:**
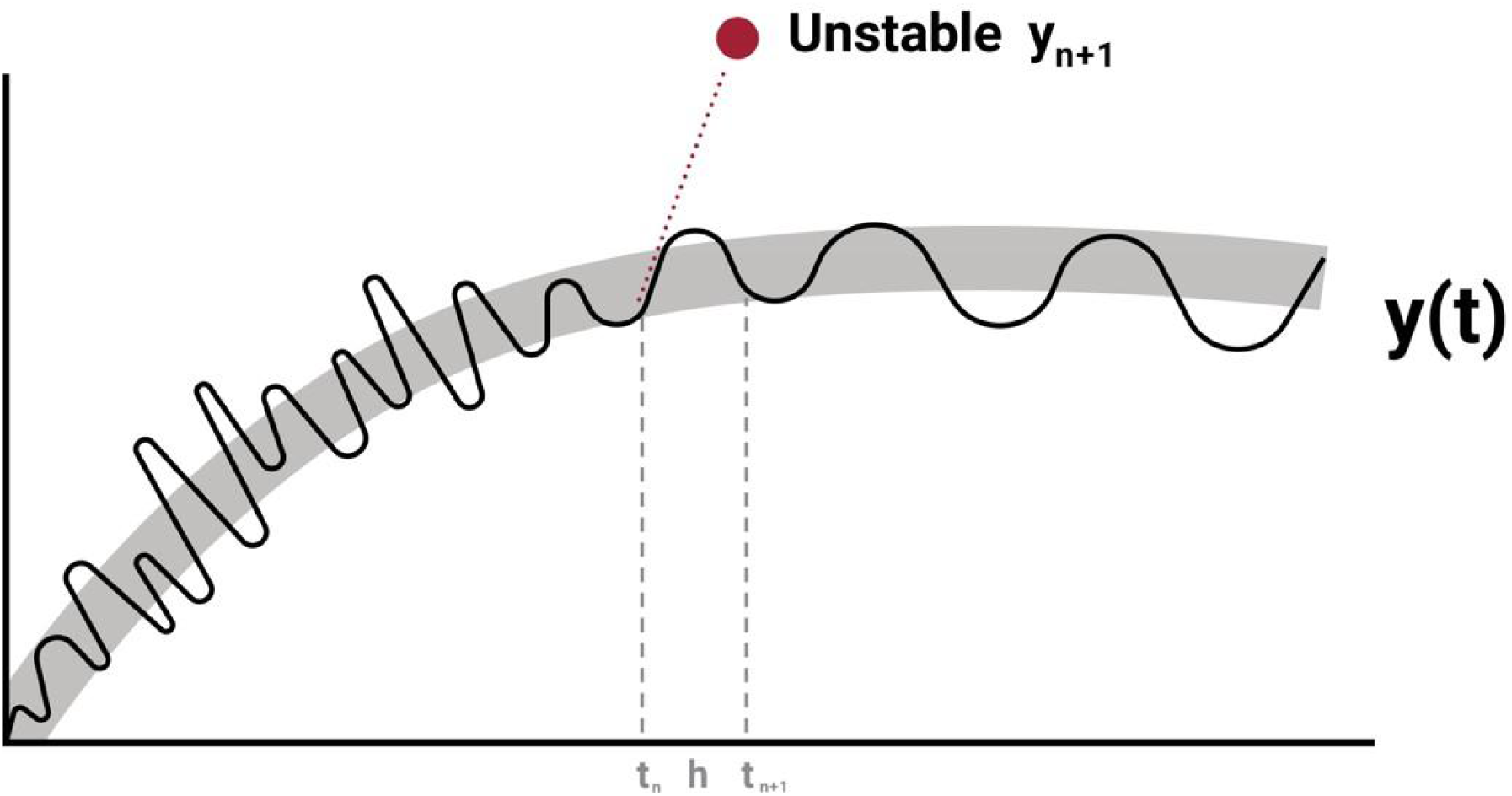
Representation of the effects of stiffness on explicit numerical predictions. Shown is a multiscale system where an explicit simulation of a stiff model is represented in dark black over the grey shade of the actual stiff ODE solution. The red line shows an extrapolation using the derivative at a value *t*_*n*_ to *t*_*n*+1_, which causes explicit Runge-Kutta methods like Euler’s method to overpredict the changes on the slow manifold manifesting in oscillations which lead to numerical instabilities. Note that the true solution may not have such oscillations and this is an amplification effect on numerical errors.

Stiffness carries over to the incorporation of QsP models into machine learning, where many common machine learning models are not well-suited for these numerically difficult equations. One way to see this is to identify the relationship between common deep learning techniques and explicit ODE solvers. Recurrent Neural Networks (RNNs) and Residual Neural Networks (ResNets) are discrete time series models which encode inductive bias of time series propagation and use current information to predict the next point in time [38,39,40,41]. Their basic form is:

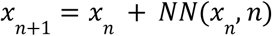

where *NN* represents a neural network. It has been noted that these architectures are equivalent under some transformations [22] and further it is known that one can generate such models as discretizations of ODEs [23,24,25]. Concretely, if one takes an ODE defined by a neural network, i.e. *x*’ = *NN*(*x*), and performs the explicit Euler discretization, one arrives at this canonical formulation of the ResNet/RNN. Similarly LSTMs can be expressed as explicit ODE discretizations with extra states for holding memory [26].

Given this direct relationship between common machine learning time series models to such explicit ODE solvers, their ability to accurately predict stiff dynamics should be suspect. Our previous work has shown this observation indeed holds numerically in other fields, where such methods tend to diverge on stiff models in cases like modeling of small chemical reaction networks to air conditioning and HVAC systems [1,2]. Here we will show that this phenomenon occurs in the case of QsP modeling by showing how a production QSP model similarly demonstrates such instability on standard machine learning methods, demonstrating the need for stiffly-aware machine learning techniques.

Given this inquiry, the clear solution would be to develop models which are trained implicitly like stiff ODE solvers. The methods for doing so come from the burgeoning field of scientific machine learning (SciML) which mixes the disciplines of mechanistic modeling with machine learning. One such an example technique was recently developed as stiff neural ordinary differential equations, where such deep learning models can be kept in their continuous form *x*’ = *NN*(*x*) and stabilized adjoint methods with stiff ODE solvers are used to fit the neural network directly within the differential equations [27]. This has been used to successfully train on data from stiff ODEs. However, this technique can take considerable compute power and currently is difficult to scale to the very large models being used in practice by QsP teams. More research into optimizing stabilized adjoint techniques will be required for these techniques to be practical across the continuum of biological models.

Another method to allow for training fully implicitly in time is the Biologically-Informed Neural Network [28,29], which is the domain translation of the Physics-Informed Neural Network (PINN) technique [19]. The core of this method is to represent the solution of a differential equation via a neural network by imposing that its derivatives satisfy the differential equation. For example, with an ODE this would be done by minimizing the loss function:

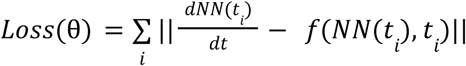

where θ are the weights of the neural network and the summation is taken over some set of time points. While these methods have shown promise in some aspects, there are some deficiencies which are the reason for the current lack of adoption in clinical and preclinical practice. For one, it can easily be shown that PINN software such as deepxde [43] are around 10,000x slower for model inference than traditional ODE solvers used in pharmacometrics software, like DifferentialEquations.jl used in Pumas1. Additionally, recent research has demonstrated that PINNs trained on stiff ODEs exhibit training difficulties and instabilities [9]. In that work it is noted that the gradient descent method:

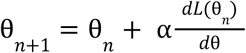

is the Euler discretization on the ODE of the gradient flow in parameter space:

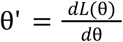

where the learning rate α is related to the Δ*t* of the Euler method. The authors show that the stiffness of this gradient flow directly correlates with the stiffness of the original model, and thus the difficulty of gradient-based optimization of PINNs increases as the model becomes more stiff. While the authors demonstrated the ability to use an adaptive explicit method as the optimizer, the test problems were PDE models with condition numbers of ∼10^5, models where explicit ODE solvers are slow but do not diverge. These “stiff” test cases have orders of magnitude less time scale separation than the >10^8 or higher condition numbers typically seen in difficult chemical reaction models. None of these previous works demonstrate the PINN training as accurate and stable on the highly stiff equations we see from QsP applications. Later in this manuscript we will show an example of PINNs failing to fully train on such examples which explains this absence.

### Continuous-Time Echo State Networks (CTESNs)

Given the noted issues with common machine learning architectures in the presence of stiffness, one might ask what the true analogue of implicit ODE solvers to deep learning architectures would be. We propose the Continuous Time Echo State Network (CTESN) as this analogue. CTESNs are a continuous-time generalization of Echo State networks, a reservoir computing method [1]. This is a scientific machine learning (SciML) which does not seek to replace mechanistic dynamical modeling with machine learning, but instead augment or combine differential equations into the machine learning architecture.

There are two alternative ways to understand the CTESN formulation. In a numerical sense, one can understand it as a neural ODE where part of the neural network (the input layer) is set to a fixed constant. Making only the last layer need to be fit is a trick which allows for the training to be fully implicit in time and generates as output a new nonlinear differential equation system that models the sub-portion of the model the practitioner was interested in. While that description provides the motivation as to how it is a transition from the previous section, an alternative explanation as a projection method against a reservoir can be more intuitive, and thus we explain the mathematics behind the architecture using this second interpretation.

In the CTESN, a randomly chosen ordinary differential equation, called a reservoir ODE, is constructed by exciting a fixed neural network layer with a chosen input whose response evolves in continuous time. Projections are then calculated from this reservoir ODE to an ensemble of solutions of the reference ODE at sampled points from the input parameter space. A map is then constructed between the input parameter space and the space of projections. A prediction at a particular test set of parameters involves constructing the projection, simulating the reservoir, and then projecting the reservoir dynamics. We call this object which can take in a set of parameters and produce a time series prediction the CTESN surrogate.

The dynamics of the reservoir ODE can be defined as:

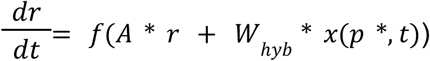

Where *x*(*p* *, *t*) is a solution at some time point in the chosen parameter space of the reference model with dimension N, *W*_*hyb*_ is a *N*_*R*_ x *N* dense random matrix that couples the physics-informed excitation to the reservoir [8], *A*is a *N*_*R*_ x *N*_*R*_ sparse random matrix representing the connections of the reservoir, and r is the current state of each of the reservoir nodes.

The projection of the reservoir’s dynamics as a response to the input can then be defined as:

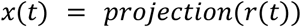

Which is the only part of the architecture that is trained and can either be linear:

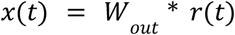

Or nonlinear [2]:

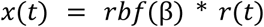

Where *W* _*out*_ is the linear projection computed by least squares using singular value decomposition (SVD), and *rbf*(β) is the nonlinear projection using a Radial Basis Function (RBF) with βas it’s coefficients. This formulation however is only for the case of surrogatizing a model with a single instance of parameters. To extend this so that the surrogate can represent outputs over a range of parameters, the projections can be reformulated slightly such that:

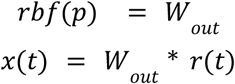

And:

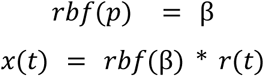

Where an RBF is used as an interpolating function that takes in model parameters as input, and outputs the weights of the projections for the linear (LPCTESN) or nonlinear case (NPCTESN). Thus, the CTESN essentially learns a method for simulating a relatively non-stiff ODE whose projects capture the original stiff ODE dynamics. Stiff ODE solves scales like O(n^3) for the number of ODEs due to internal LU-factorizations. Compared to the O(n) scaling of non-stiff ODE solvers, this suggests that such a transformation to a non-stiff ODE projector form would give an acceleration that would grow as the systems become larger and more complex, a result which we will demonstrate in the preceding sections.

Because the CTESN projects are trained via implicit linear solves (SVD factorizations) instead of gradient-based optimization, this avoids the previously mentioned gradient pathologies that arise in other surrogate methods such as PINNs and LSTMs [9]. This formulation of the CTESN also takes advantage of the heuristics and knowledge embedded within adaptive time steppers for stiff differential equations, automatically choosing key points in the reference model’s trajectory in a way that gravitates towards the more difficult dynamics. This allows the training data to capture the reference model’s evolution over multiple time scales which would be difficult using fixed step discrete schemes [1]. These underlying numerical properties are the reasons we propose the CTESN’s to model very stiff systems.

### Case Studies

#### CTESNs Handle a Leucine Model where Traditional Deep Learning Fails

We first consider a model capturing the metabolism of the Leucine amino acid in humans [3,36]. While this model is not sufficiently large enough to demonstrate acceleration by surrogate techniques, it was chosen since it was previously noticed to be a model with many difficult stiff behaviors, causing potential issues with many numerical ODE solvers, and thus would serve as a benchmark for the ability for surrogate techniques to accurately capture dynamics which widely vary over large parameter spaces.

We construct a surrogate of this model with an input parameter range of +/-50% of the base parameter set. We sample 100 sets of parameters from this space using Latin hypercube sampling, and simulate the model at each parameter. We then compute projections to each of these solutions from a reservoir system, and use a radial basis function to learn a nonlinear map from the input parameter space to the space of projections. The reservoir system uses a weight matrix with negative eigenvalues and a softplus activation function.

We compare the CTESN prediction to other common machine learning methods to learn dynamical systems, namely the PINNs and LSTMs. The PINNs network architecture consisted of 3 fully connected hidden layers with 128 hidden nodes and a softplus activation function, and was trained for 2000 epochs using the ADAM optimizer. The LSTM architecture consisted of 3 LSTM layers with 128 hidden nodes and was trained for 2000 epochs with the ADAM optimizer. Both the PINNs and the LSTMs were trained by generating time-state pairs throughout the timespan at timepoints informed by the ODE solver. Figure 2 shows the surrogate prediction at output dimensions 5 and 7 on unseen parameters. In addition to superior predictive performance, the CTESN predicts reasonably well over an entire space in contrast to the PINN and LSTM implementation. The prediction error from the CTESN can be understood by examining Figure 3, which shows the variation of output dimension 5 throughout the parameter space. This lends insight into why the predictive error in output dimension 5 is higher than in 7, which is better behaved.

**Figure 2:**
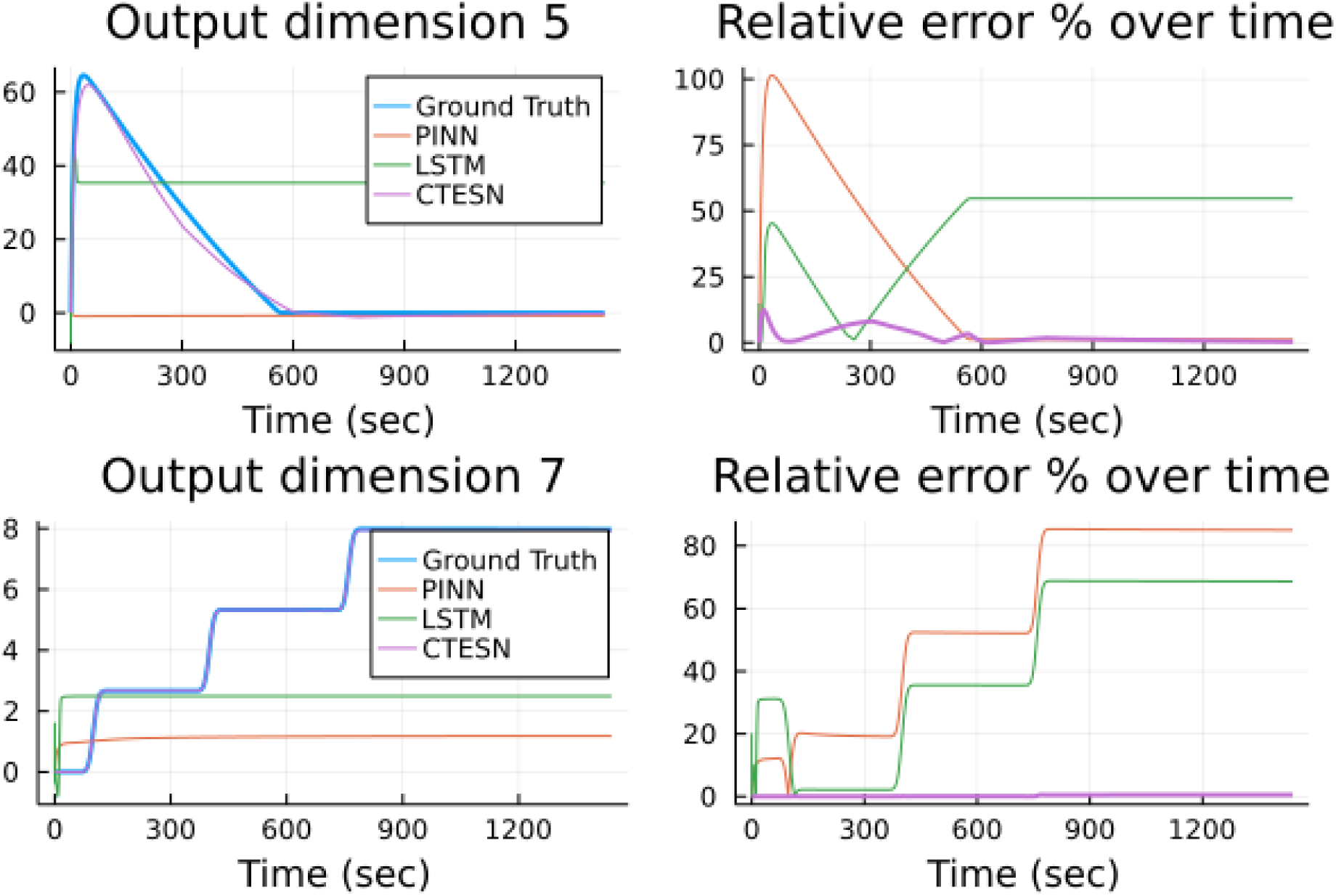
Performance of traditional scientific machine learning models vs the CTESN on the stiff Leucine model. The ground truth is given in blue while the predictions from the PINN, LSTM and CTESN are in red, green and purple. The LSTM was trained with 3 hidden LSTM layers of length 128 and was run for 2000 epochs and the PINNs used 3 hidden fully connected layers of length 128, and were trained on pairs of time and state vectors. The CTESN was trained by calculating nonlinear projections (NPCTESN) from a reservoir of size 14 to 100 time series in the chosen input space, and . The error from the CTESN can be explained by the variation of the output dimension 5 in the chosen input space, as shown in Figure 3.

**Figure 3:**
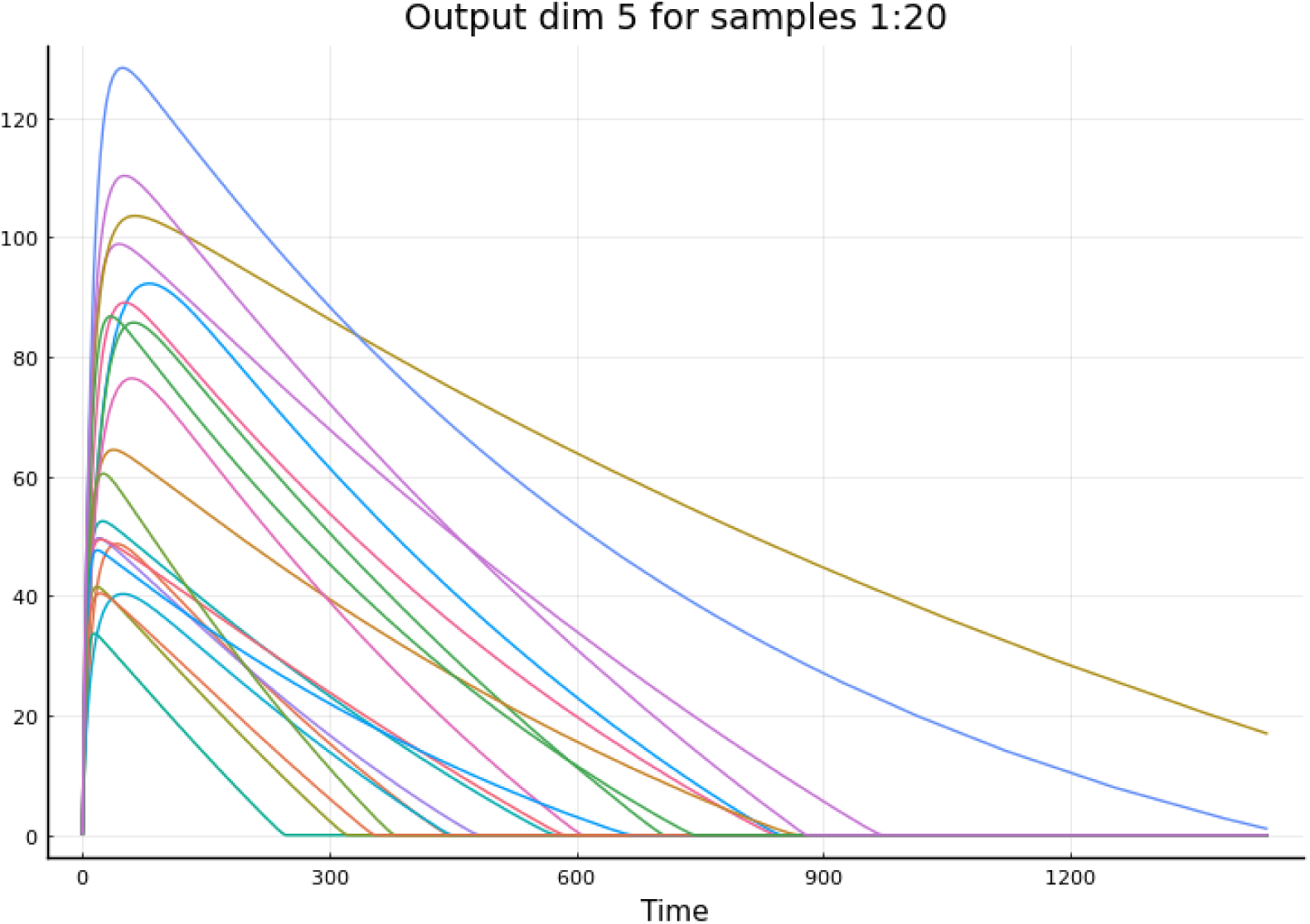
Variation of output dimension 5 across chosen input space. The ODE is simulated at 100 different points in the input parameter space, obtained via Latin Hypercube sampling. Output dimension 5 was plotted for 20 such points. The large variation in behaviour across the space affects the quality of the surrogate model predictions in such a way that only the CTESN is able to achieve relative accuracy.

#### CTESNs Reliably Accelerate Large QSP Models from SBML

In order to make it easy to test CTESN surrogate generation on large biochemical reaction network models, we created biological model importers for the ModelingToolkit.jl [32] symbolic language for the common CellML, BioNetGen, and SBML file formats. This allows for automatically importing models from repositories such as the CellML Physiome Project [31], the SBML Biomodels Database [33], and more to all be simulated with Julia’s SciML tools [34] and be automatically surrogatized via CTESNs.

Using this infrastructure, we tested the CTESN automated surrogatization procedure on a WUSCHEL gene model [10] from the SBML Biomodels Database. While this model is not specifically a QsP model, it is a relatively large ODE model (1265 ODEs and 522 input parameters) with structures such as mass action kinetics, Michaelis-Menton, and Hill function interactions which are indicative of the common forms seen in QsP models. The model predicts the dynamical reorganization seen in experiments where cells including the WUSCHEL domain, are ablated, and it also predicts the spatial expansion of the WUSCHEL domain resulting from removal of the CLAVATA3 signal. This model is noted as stiff since implicit methods were required to adequately numerically solve the equations.

One advantage of such surrogate models is that it can be used to generate nonlinear dynamical systems which are fast and only require predicting the biomarkers of interest. To simulate this process, we trained a CTESN on the WUSCHEL model surrogatized with respect to 10 parameters predicting 10 outputs (the first 10 reacting species) over a time span of 0 to 20. We sampled on +/-10% of base parameters (due to numerical instability of ODE solvers beyond this range). We sample 50 sets of parameters from this space using Latin Hypercube sampling and the model and compute 50 nonlinear projections from the reservoir ODE to the reference model. We then interpolate through these projections using a radial basis function as described in the previous section.

As a baseline, the MATLAB SimBiology took 0.7097 seconds to simulate the model. The baseline Julia simulation with DifferentialEquations.jl took 0.005091 seconds to simulate using CVODE BDF method [11], while the surrogate took 0.0009801seconds to predict the same time series with a relative error of less than 1% across all the 10 predicted outputs^2^. This amounts to a **5.2x speedup** against the Julia differential equation solvers and a **724x speedup** against the MATLAB SBML tools. The training procedure can be parallelized to the cost one ODE solve.

The predictions are shown in Figure 4, and Table 1 shows the relative error of each output. Since this was performed using the JuliaSim default CTESN surrogate training hyperparameters, this demonstrates that the training process is reasonably stable without tuning the training process to new problems, allowing for arbitrary chemical reaction network models from such file formats to be quickly surrogatized into ML-accelerated forms for further analysis, such as global sensitivity analysis and virtual population generation. Figure 5 demonstrates that this acceleration increases for models simulated over longer time spans.

**Figure 4:**
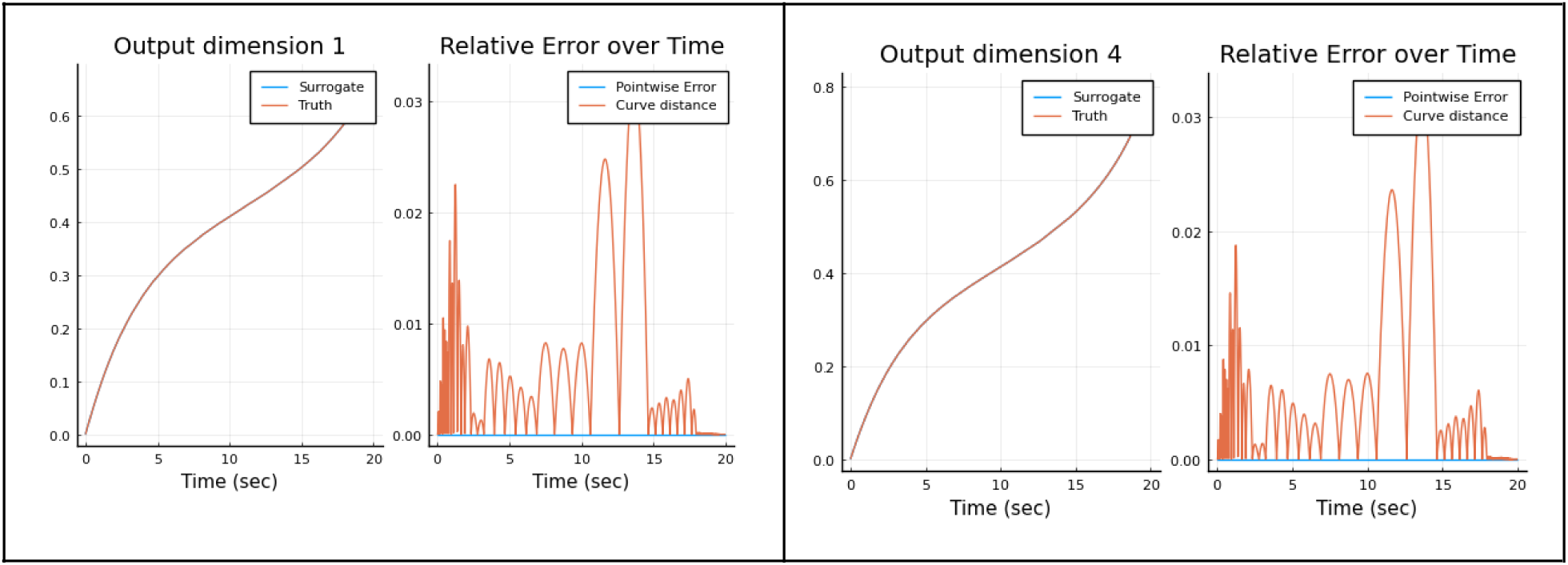
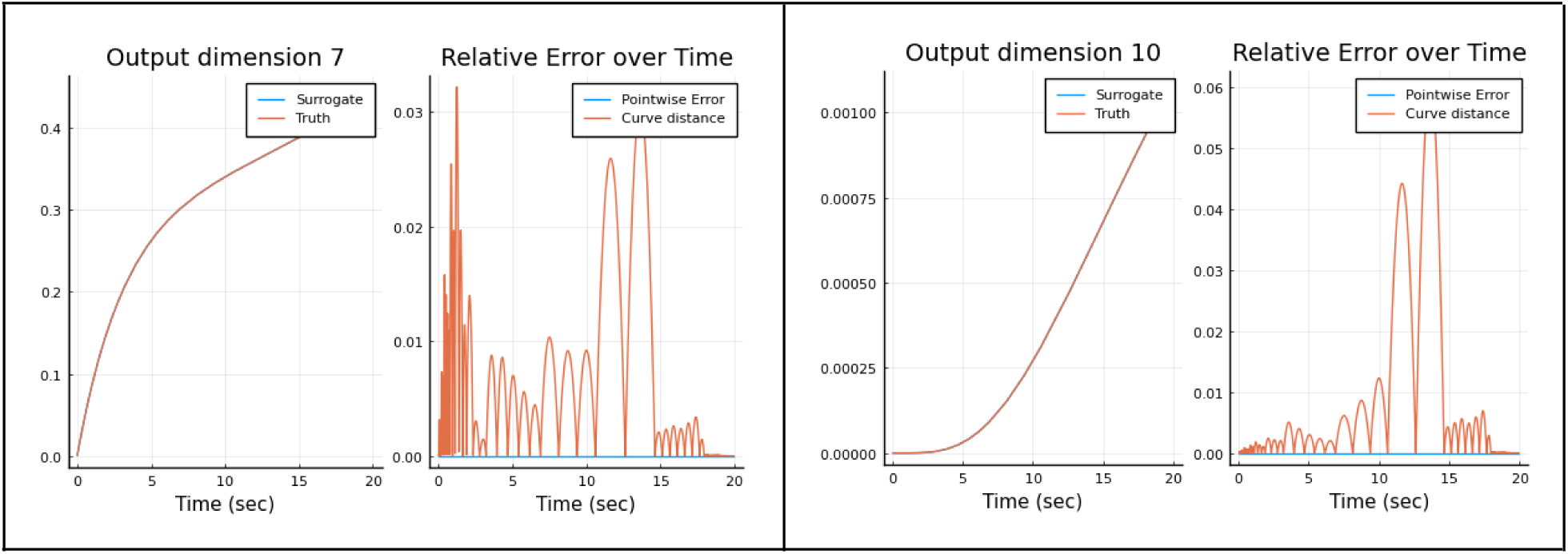
CTESN prediction of the WUSCHEL model. Each cell represents the prediction of some 4 chosen outputs. On the left of each cell, the blue line represents the prediction from the surrogate and the red represents the reference test solution (ground truth). On the right of each cell, two error metrics are plotted: the blue represents pointwise relative error percentage for each sampled prediction and the blue line represents a curve distance, which is the minimum distance from a prediction to any ground truth point along the trajectory.

**Table 1:** Relative error percentages of each output from the CTESN compared with a reference test solution.

| No. | Output | Error (%) | No. | Output | Error (%) |
| --- | --- | --- | --- | --- | --- |
| 1 | Brusselator Variable A in Cell 17 | 0.1403 | 6 | Wuschel in Cell 103 | 0.0385 |
| 2 | Brusselator Variable A in Cell 67 | 0.1248 | 7 | Brusselator Variable A in Cell 42 | 0.1352 |
| 3 | Brusselator Variable A in Cell 176 | 0.1221 | 8 | Brusselator Variable B in Cell 249 | 0.0839 |
| 4 | Brusselator Variable A in Cell 99 | 0.0728 | 9 | Layer 1 Specific Protein in Cell 182 | 0.0057 |
| 5 | Layer 1 Specific Protein in Cell 182 | 0.000 | 10 | Diffusible Wuschel Inhibitor in Cell 88 | 0.1336 |

**Figure 5:**
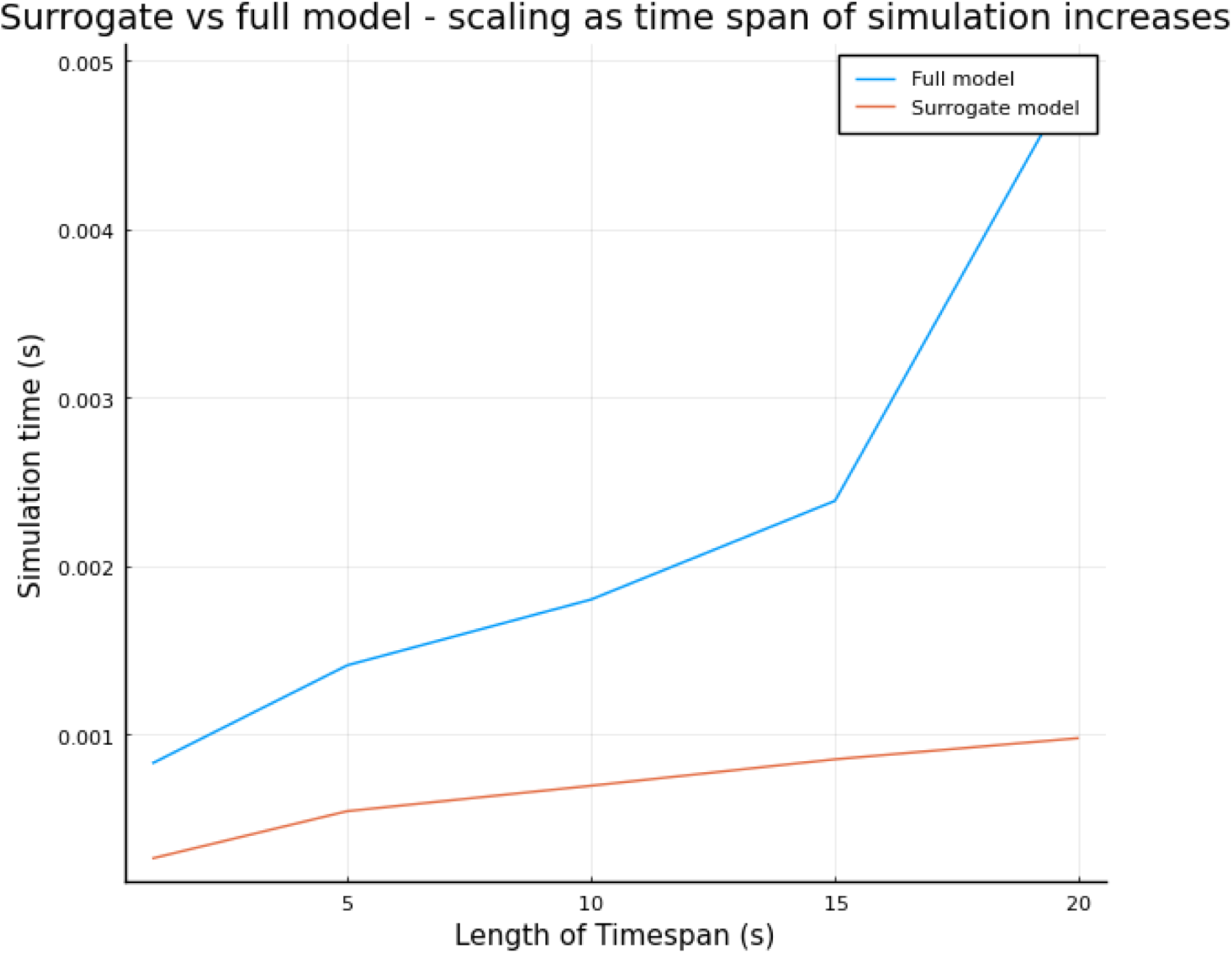
Scalability of the CTESN surrogate with respect to time span. Shown is the time for simulation with the trained surrogate on time spans from 0 to x, where x is shown as the x-axis of the plot. The simulation time is shown in the y-axis in seconds. The surrogate scales linearly throughout.

#### CTESNs Acceleration Grows with Model Size on Biopharmaceutical Partial Differential Equations

To test how the CTESN surrogate scales for models of varying sizes, we used semi-discretizations of a reaction-diffusion equation which is a classic model for spatially-varying chemical reaction systems. We chose a PDE model of a reaction including three species, *A, B, C*, with parameters *D*, α _1_, α_2_, α _3_, β_1_, β_2_, β _3_. The PDE followed the form:

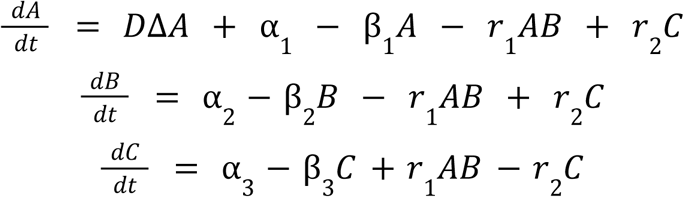

With Neumann boundary conditions:

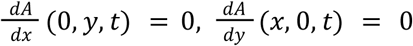

We solved this on the domain [0, 1] × [0, 1]discretized into *N*points along each axis (giving *N*^2^ knots per timestep), with zero initial conditions, for *N ∈* {32, 50, 75, 100, 125, 150, 175, 200, 225, 250}, giving our largest solution (3)(250^2^) = 187500 data points per timestep with time spanning to *t* = 100^3^. One use-case of this model might be if the user was interested in the average concentration across the grid of each species, the concentrations of just one species, or the concentration of the species at a point. However, due to the Laplacian in the first species’ equation, this equation can’t be solved at just one point or a particularly coarse grid of points. Therefore, a surrogate might be necessary so that, in the optimal case, a scientist could glean information about a particular point without having to dedicate the resources to simulating the entire PDE.

In this example, the target solutions were near the parameters *D* = 100, α *_k_* = 1, β *_k_* = 1, *r*_*_j_*_ = 1for *k* = 1, 2, 3 and *j* = 1, 2 and in the outputs corresponding to the grid {0, 1/4, 1/3, 1/2, 2/3, 3/4, 1} × {0, 1/4, 1/3, 1/2, 2/3, 3/4, 1}. However, this model shows a scalability of surrogates for PDE applications where solving several full solutions at various points in a parameter space is difficult or impossible. The evaluation time of the surrogate model theoretically should only scale corresponding to the size of the output grid (in this case 3(7 ^2^) = 147), which is independent of *N*.

To surrogatize the model, 100 sets of parameters were randomly chosen (using Latin Hypercube Sampling) from ± 10% of the target values, and the model was fully solved for these sets of parameters. Then, a nonlinear projection CTESN (NPCTESN) was trained on these 100 samples for each given *N*, with each surrogate employing a reservoir size of 10. In Figure 6a below, we see that evaluating the surrogates produced for each *N*grows minimally, from 0. 024692seconds to 0. 377seconds, where the full model grows quadratically from 0. 024 seconds to 21. 405seconds. This represents a relative **acceleration of 56x** for the largest model. The surrogate models are able to approximate the whole time series with a relative error of below 5% in all cases while generally increasing in accuracy as the discretization becomes increasingly fine, as seen in Figure 6b.

**Figure 6a:**
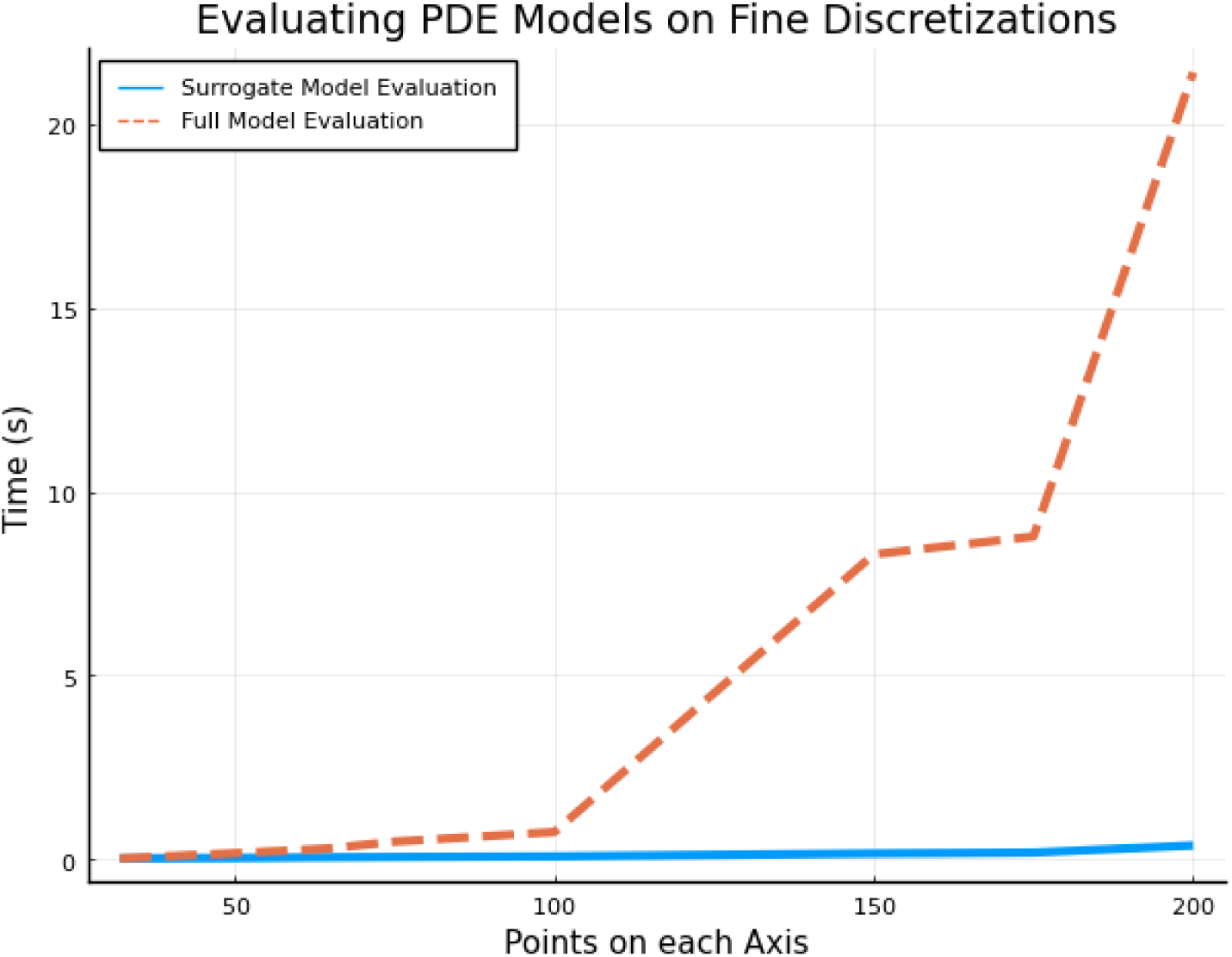
Performance of surrogate vs full order model on finer discretizations of a reaction diffusion PDE. The blue line indicates the time taken for the surrogate to predict an output time series from the PDE while the red line denotes full PDE simulation time.

Each surrogate model in this example was trained on 100 examples of the full solution of the PDE, each solved using parameter values within 10% of the target parameters, then reducing the output to the reduced spatial grid. It was found that these surrogate solutions were generally accurate overall to within 5% error, the error found was highly dependent on the choice of parameters of the full solution that the surrogate was trained on. To track this idea, the seed of the random number generator was set to a fixed value for training and, as seen in Figure 6b, the surrogates track closely together for the same sets of parameters.

The variability seen in Figure 6b for random *p* clearly vanishes once the parameter samples are fixed. This variability is largely attributable to such a high dimensionality of the parameter space, as the PDE is parameterized by nine values. However, the CTESN approach gives a realistic estimation of a subset of grid points without the need to simulate every grid point in the domain for every variable.

**Figure 6b:**
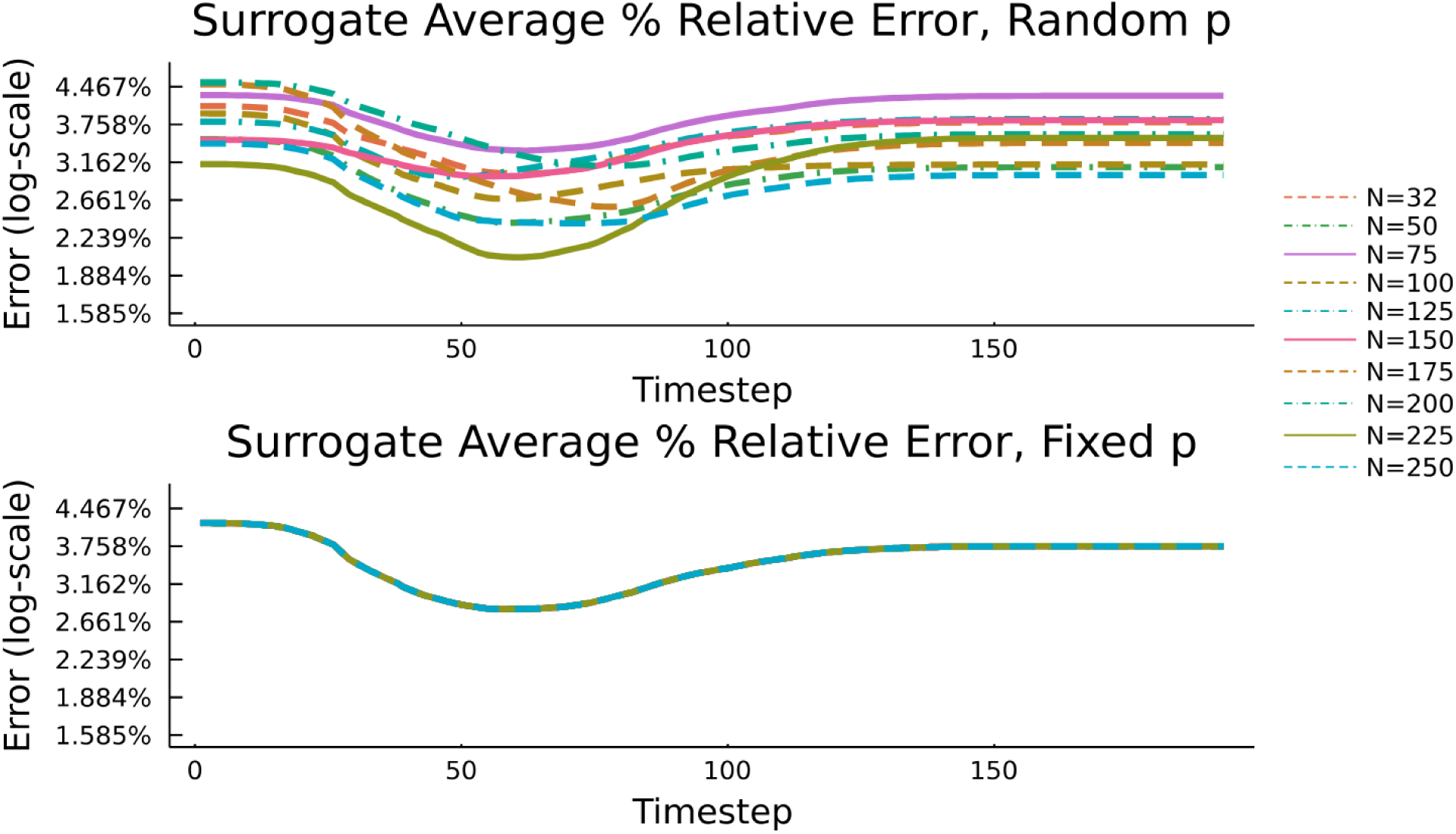
Average % relative error of surrogate from its respective full discretization on output points over each timestep for 15 solutions within 10% of the target parameters. (Top) Surrogates trained on random *p* for each *N*. (Bottom) Surrogates trained on fixed *p*for all *N*

#### Conclusions and Future Work

Quantitative pharmacology has always proposed major methodological challenges which needed to be addressed in order for the field to ensure accurate predictions. With clinical pharmacology, the famous work of Beal and Shiner was the catalyst that allowed for nonlinear mixed effects modeling to become common practice in trial analysis. With quantitative systems pharmacology, the foundation is decades of research in methods for stiff ODE solvers which generally used such chemical reaction networks as the hard examples [44,45,46,47]. Now with the rise of machine learning, we are demonstrating that it will be no different from before: pharmacology can take as influence the standard methods from machine learning but will need to adapt them in order for them to reach the stability requirements required for handling such large-scale and stiff biochemical reaction models.

In this manuscript we described how the properties of QsP models would give many machine learning techniques issues, and demonstrated their inability to capture the dynamics of a production QsP model with known numerical difficulties. However, motivated by a theoretical backing, we hypothesized that the CTESN corresponds to the “stiff ODE solver of machine learning”, and demonstrated its ability to handle the ill-conditioned problems imported from SBML model databases and generated from reaction-diffusion PDE discretizations.

While machine learning architectures are typically difficult to train, the robustness and implicitness of this technique makes it amenable to many QsP workflows. In all of the examples, we restricted ourselves to no more than 100 training points for the training of the emulators, and note that the training time (i.e. the SVD-factorization) was dwarfed by this sampling procedure. Thus with roughly the real-time budget of one ODE solve, 100 simultaneous ODE solvers can be distributed over a cloud compute architecture to build this surrogate for researchers to continue with virtual population and global sensitivity analyses. This is a major relative advantage over surrogate techniques which require gradient-based optimization, a very costly and mostly serial training procedure which in many cases takes longer than the requested analyses themselves. In fact, all of our experiments with the CTESN were able to be run on simple laptops without requiring any GPU acceleration.

As such, this manuscript demonstrates that the recent scientific machine learning (SciML) techniques which seek to integrate the techniques of machine learning with mechanistic modeling may provide a realistic path forward for integration of machine learning into standard QsP workflows. This method does not attempt to replace differential equation modeling with neural networks, but instead is an architecture which defines differential equations in part by neural networks and fuses the properties of stiff ODE solvers into it, making it very similar to the kinds of QsP models it is trying to emulate and thus improving its predictive performance. This is a demonstration of just one possible technique at the intersection between these two distinct modeling disciplines and points to such hybrid approaches as being potentially fruitful avenues of future research.

## Acknowledgement

The information, data, or work presented herein was funded in part by the Advanced Research Projects Agency-Energy (ARPA-E), U.S. Department of Energy, under Award Numbers DE-AR0001222 and DE-AR0001211, and NSF awards OAC-1835443 and IIP-1938400. The views and opinions of authors expressed herein do not necessarily state or reflect those of the United States Government or any agency thereof.

## Footnotes

1 A demonstration of this 10,000x performance difference is documented at https://github.com/SciML/DataDrivenDiffEq.jl/issues/192#issuecomment-770456859

2 Details of the SBML reader are still being checked and timings may change in the next revision.

3 For the particular model chosen, arbitrarily fine discretizations are actually not necessary. At our target parameter values and outputs, the discretized full model for N=32 had relative error that was numerically zero.

